# Aboveground fungal lethal effects of local plants on invasive plant *Ageratina adenophora* seedlings

**DOI:** 10.1101/2025.01.21.634076

**Authors:** Zhao-Ying Zeng, Zi-Qing Liu, Ai-Ling Yang, Yu-Xuan Li, Yong-Lan Wang, Evan Siemann, Bo Li, Han-Bo Zhang

## Abstract

Native community resistance to plant invasion may reflect microbial effects on invader establishment. Here, we compared the effects of the aboveground and belowground microbes and leaf functional traits of 25 native plant species on the germination and survival of invader *Ageratina adenophora*. Results show that leaf physical and chemical traits vary among plant species and shape the aboveground microbial community. Local plants more phylogenetically close to *A*. *adenophora* harbour more similar pathogen communities to those *A. adenophora.* The aboveground tissue inoculations had more adverse effects on *A. adenophora* establishment by delaying germination time and decreasing the germination rate and seedling survival than did soil inoculations; aboveground tissues from local plants, which are more closely related to *A. adenophora*, cause longer germination times and greater seedling mortality of *A. adenophora*. Moreover, local plant aboveground fungi, rather than bacteria, adversely impact seedling survival, and most of the detrimental fungal pathogen ASVs are specialists. Several strains isolated from dead seedlings caused by aboveground tissue inoculations belonging to *Alternaria*, *Fusarium* and *Stagonosporopsis* were verified to kill more than 60% of *A. adenophora* seedlings and thus have great potential as biocontrol agents for *A. adenophora*. This is the first study to provide evidence for the effects of phylogenetic relatedness and leaf traits (both functional and microbial) on local community resistance to invaders through the regulation of germination and seedling survival.

## Introduction

In many cases, local plant community resistance to invader is determined primarily by phylogenetic relatedness and functional trait similarity between resident plants and invaders (Li et al., 2015; van Kleunen et al., 2018; Zheng et al., 2018). Plant phylogenetic relatedness reflects the similarity of functional traits of plant species, including physical and chemical traits, as well as the microbial community inhabiting above and belowground. For example, phylogenetic relatedness and phenotypic trait similarity (including biomass, specific leaf area, leaf dry matter content and root traits) are positively associated (Fitzpatrick et al., 2017). Plant leaf traits such as leaf mass per area and leaf nitrogen and phosphorous concentrations are significantly correlated with microbial community structure; moreover, the increasing phylogenetic distance among plant species leads to more dissimilar bacterial communities in the rhizosphere or phyllosphere (Emmett et al., 2017; Kembel and Mueller, 2014; Kembel et al., 2014). These interactions in turn contribute to invasion of phylogenetically distant species via the regulation of plant growth. For example, Parker et al. (2015) reported that phylogenetically distinct species are more likely to become invasive because of their lower susceptibility to local pathogens.

In addition to host growth, phylogenetic and functional relatedness also affect seed germination and seedling mortality, key processes in determining plant community composition and dynamics (Huang et al., 2016; Kimball et al., 2010). For example, Liu et al. (2012) reported that negative effects on seedling survival decreased with increasing phylogenetic distance of neighboring trees due to weaker soil pathogen effects. By contrast, studies showed that phylogenetically close species had more similar germination niche and survival patterns, because they required similar soil condition (including nutrients, moisture, texture and mutualist microbes) (Burns and Strauss, 2011), thus invader may better to achieve germination and survival when they introduce close related community. Nonetheless, there have been no experiments to test whether the phylogenetic and functional relatedness of local resident plant communities to invaders plays a role in their resistance to invasion via aboveground (mainly leaf tissues) microbial community effects on invader’s germination and seedling survival.

Leaf litter in natural systems is important for plant population establishment and community dynamics (Jessen et al., 2023; Lamb, 2008; Xiong and Nilsson, 1999). However, research to date has focused mainly on the physical (e.g., maintaining soil moisture and temperature, increasing nutrients and reducing light) or chemical (e.g., releasing allelochemicals) effects of litter on seedling emergence and growth (Demey et al., 2013; Jessen et al., 2023; Möhler et al., 2018; Zhang et al., 2017). Recent studies have indicated that foliar microbes could have more negative effects on plant growth than soil microbes due to relatively high pathogen levels in the phyllosphere (Fang et al., 2019; He et al., 2023; Whitaker et al., 2017). Moreover, studies have shown that closely related plants share more foliar pathogens or are more susceptible to the same foliar pathogen (Gilbert et al., 2012; Gilbert and Webb, 2007). In particular, plant pathogens inhabiting senescing leaves could be sources of microbes that negatively impact seedlings and thus regulate their density (Márquez et al., 2011; Tanunchai et al., 2022). Here, we hypothesized that local plant aboveground microbes can act as more effective weapons than soil microbes that limit invader’s seed germination and seedling survival, depending on the phylogenetic relatedness and functional trait similarity between resident and alien plants.

*Ageratina adenophora* (Sprengel) R.M. King & H. Robinson (Asteraceae) is one of the most common invasive weeds worldwide, especially in China (Gu et al., 2021; Wang and Wang, 2006). Recently, Fang et al. (2019) and Zeng et al. (2024) reported microbes in the aboveground tissues of *A. adenophora* adversely affects seed germination and seedling survival;. Chen et al. (2020a) and Zeng et al. (2023) reported that fungal cross-transmission between *A. adenophora* and surrounding plants is common. In this study, we quantitatively compared the effects of aboveground tissues and belowground soils collected from 25 local native plants on *A. adenophora* seed germination and seedling survival under controlled experimental conditions. We also measured the aboveground and belowground microbes and leaf functional traits to understand local plant physicochemical and microbial effects on *A. adenophora* seed germination and seedling survival. We predicted that 1) aboveground microbes of local plants would reduce *A. adenophora* seed germination and seedling survival more strongly than would soil microbes because aboveground tissue would have a greater proportion of pathogen abundance than would soil. 2) Plants that are more closely related to the invader *A. adenophora* will present more similar aboveground pathogen communities to those in *A. adenophora*; thus, aboveground tissues of plants more closely related to *A. adenophora* will have greater negative effects on its seed germination and seedling survival.

## Results

### 1. Aboveground tissues had more adverse effects on seed germination and seedling survival than did belowground soils

At G0 inoculation, all three factors, inoculum sample type (soil or plant tissue) and plant species identity and sterilization significantly affected seedling survival (SSR_G0) (all *P*<0.05). Sample type and plant species identity significantly affected germination time (GT) and germination rate (GR) (both *P*<0.05), but sterilization did not (*P*>0.05, Table S2a). At G21 inoculation, plant species and sterilization significantly affected the seedling survival rate (*P*<0.05, Table S2b). Specifically, most local plant tissue inoculations had more adverse effects on *A. adenophora* germination time, germination rate and survival rate than soil inoculations (*P*<0.05, Fig. 1&S2). Compared with the control (plant tissue-free and soil-free), the aboveground tissues of all 26 species (regardless of whether they were unsterilized or sterile), but the soils of only two species (unsterilized *Vicia sepium* and sterile *Capparis bodinieri*) significantly delayed the germination time (all *P*<0.05); similarly, the aboveground tissues of nineteen species but the soils of only four species (regardless of whether they were unsterilized or sterile) significantly decreased germination rate (all *P*<0.05, Fig. S2).

**Figure 1.**
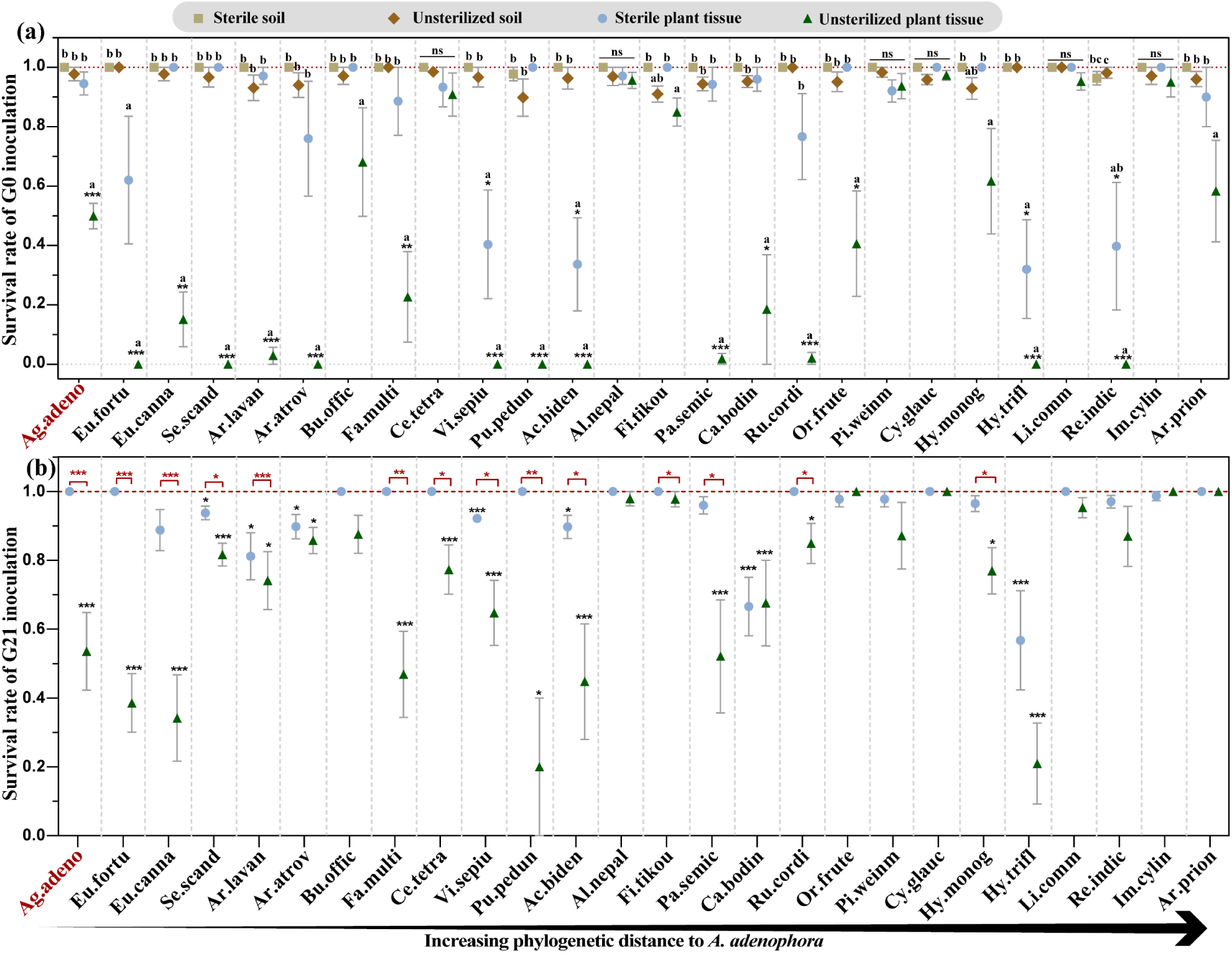
Effects of plant aboveground tissues or belowground soil inoculation on the *A. adenophora* seedling survival. The effects of sterile and unsterilized aboveground tissue or belowground soil inoculation of the 26 species on the *A. adenophora* seedling survival rate after G0 inoculation (a) and the effects of sterile and unsterilized aboveground tissue inoculation of the 26 species on the *A. adenophora* seedling survival rate after G21 inoculation (b). The black * represents a significant difference compared with the control via Nonparametric Mann-Whitney U tests. Different lowercase letters indicate significant differences among the four treatment groups of one species in panel (a) via Kruskal-Wallis test. Red * represents a significant difference between the nonsterile and sterile groups in panel b via Nonparametric Mann-Whitney U tests. The “ns” means nonsignificant. The red dotted line represents the average value for the control treatment (plant tissue-free and soil-free treatment), which was 1.00±0.00. * *P* < 0.05, ** *P* < 0.01, *** *P* < 0.001. Means ± 1 SE, n = 5. The abbreviations used for the abscissa are listed in Table S1, and the red text highlights exotic *A. adenophora*.

In terms of seedling survival, the aboveground plant tissues of sixteen species (unsterilized) and four species (sterile) presented significantly lower seedling survival rates than did those of the control (plant tissue-free and soil-free), of which the unsterilized plant aboveground tissues of eight species caused all seedlings died, and the unsterilized plant tissue inoculations of fifteen species caused significantly lower survival rates than sterile tissues did at G0 inoculation (all *P*<0.05, Fig. 1a). Similarly, the plant aboveground tissues of sixteen species (unsterilized) and seven species (sterile) resulted in a significantly lower seedling survival rate than did the control, of which the unsterilized plant tissue inoculations of fourteen species resulted in a significantly lower than sterile plant tissues at G21 inoculation (all *P*<0.05, Fig. 1b). Surprisingly, the seedling survival rate did not significantly differ among the unsterilized, sterile soil inoculations and control (plant tissue-free or soil-free) across the 26 species (all *P*>0.05, Fig. 1a).

### 2. Relationships among plant phylogenetic relatedness, leaf functional traits and microbial communities

Both the bacterial and fungal communities significantly differed across the 26 plant species and significantly differed between aboveground tissues and soils (R^2^: 0.149∼0.266, all *P* < 0.05; Fig. 2a-b). In particular, the predictive bacterial and fungal plant pathogens presented greater relative abundances in aboveground tissues than in soils (all *P* < 0.05, Fig. 2c-d). Leaf physical and chemical traits also differed among the 26 plant species (R^2^ = 0.595, *P* = 0.001 for physical; R^2^ = 0.915, *P* = 0.001 for chemical, Fig. S3a-b). Nonetheless, plant genetic distance only accounted for aboveground bacterial community dissimilarity (r*_Mantel_* = 0.34, *P* = 0.018); for functional traits, leaf physical dissimilarity was positively correlated with aboveground bacterial community dissimilarity (r*_Mantel_* = 0.26, *P* = 0.001), and leaf chemical dissimilarity was positively correlated with aboveground fungal community dissimilarity (r*_Mantel_* = 0.15, *P* = 0.014) (Table S3).

**Figure 2.**
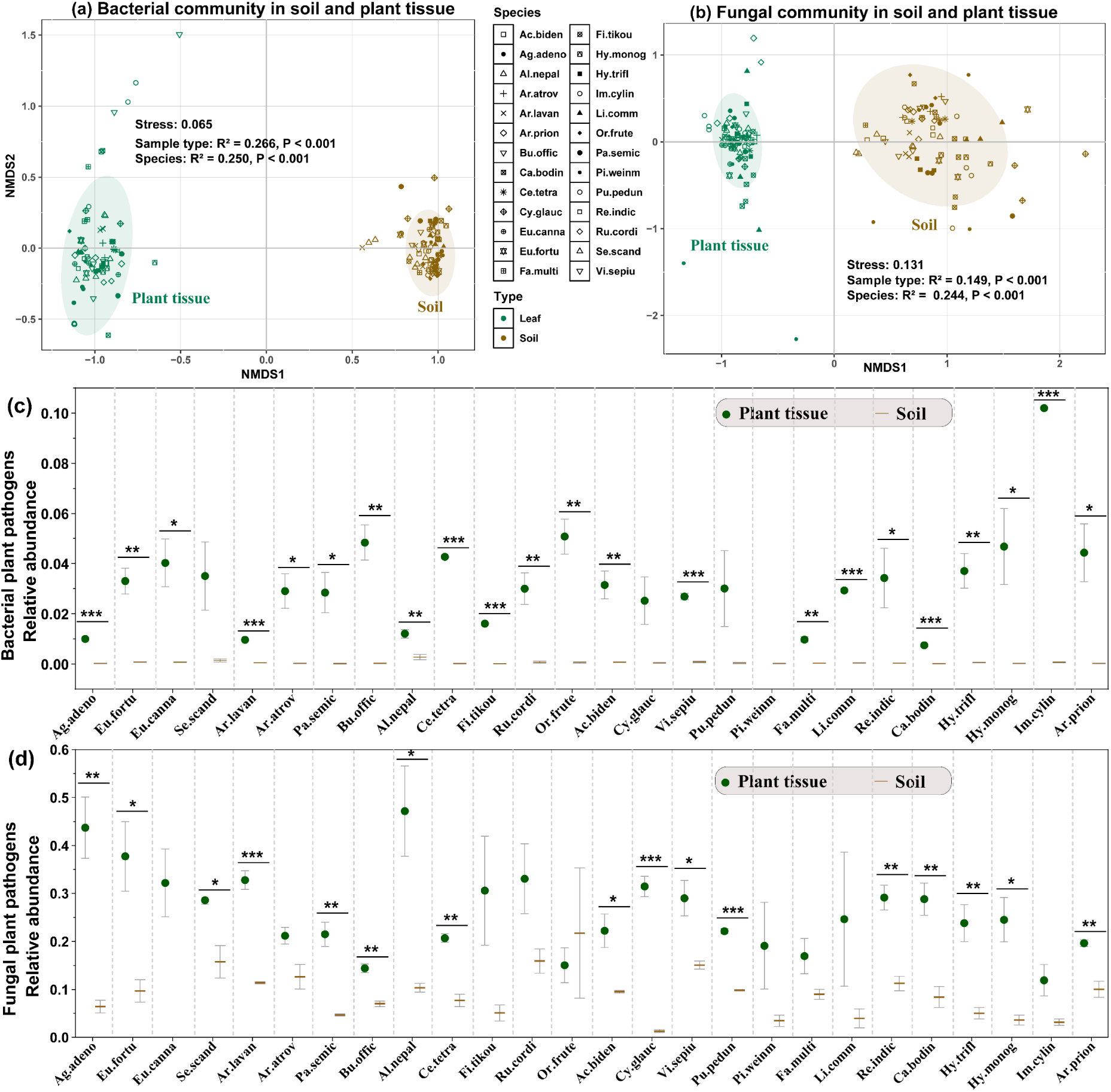
Microbial community structure and plant pathogens in the inoculum aboveground tissues and belowground soils. Bacterial (a) and fungal (b) community structure in the aboveground tissues and soils across 26 plant species are shown on the basis of Nonmetric multidimensional scaling (NMDS) ordinations of Bray–Curtis dissimilarity matrices. Different colour points indicate different sample types: green represents aboveground tissues, and brown represents belowground soil. Different shapes indicate different plant species. R^2^ represents explained variation by group, and *P*<0.05 represents significant dissimilarity among groups. Relative abundances of bacterial (c) and fungal (d) plant pathogens in the aboveground tissues and belowground soils are presented. Nonparametric Mann–Whitney U tests were used to calculate significant difference in relative abundance of plant pathogens between aboveground tissues and belowground soil from one plant species, * *P* < 0.05, ** *P* < 0.01, *** *P* < 0.001. Means ± 1 SE, n = 3. The abbreviations used for the abscissa are listed in Table S1.

### 3. Factors associated with local plant leaves account for the effects of aboveground tissue inoculation on seed germination and seedling survival

Due to aboveground tissue from most plant species caused adverse effects on *A. adenophora* seed germination and seedling survival, while soils from three plant species at most caused adverse effects on *A. adenophora* seed germination (see Fig. 1&S2), we further identified factors associated with local plant leaves that account for the effects of aboveground tissue inoculation on germination and seedling survival. Under unsterilized plant aboveground tissue inoculation, the phylogenetic distance from local native plants to *A. adenophora* was positively correlated with species effects on *A. adenophora* germination time (R^2^=0.292, *P*=0.005; Fig. 3a) and negatively correlated with the species effect on the *adenophora* seedling survival rate of G0 inoculations (R^2^=0.169, *P*=0.042) but not with those of G21 inoculations (*P*>0.05) (Fig. 3c). With sterile plant tissue inoculation, phylogenetic distance was negatively correlated with only the germination rate (R^2^=0.191, *P*=0.029; Fig. 3e).

**Figure 3.**
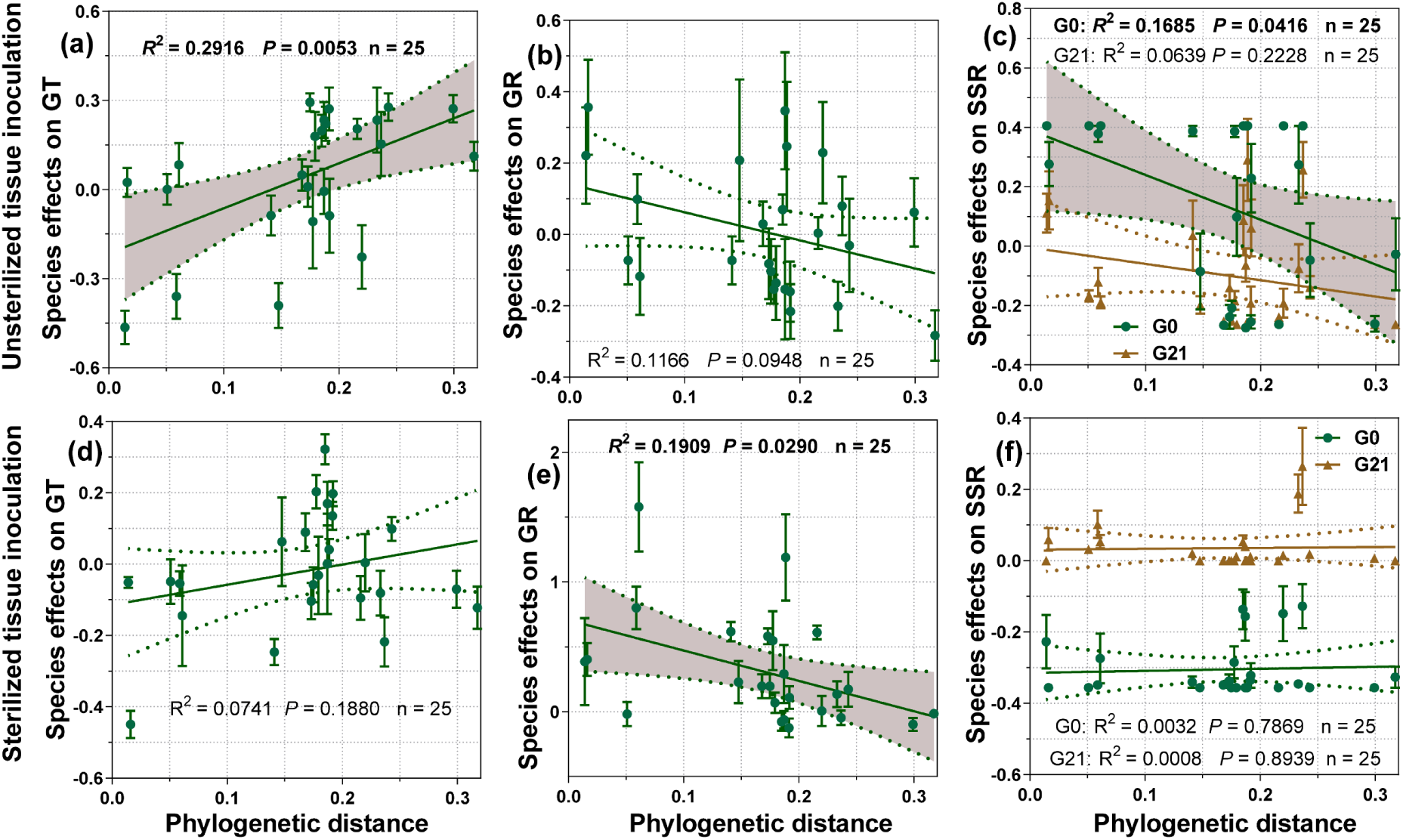
Relationships of phylogenetic relatedness with seed germination and seedling survival. The linear relationships of species effects (SE) of unsterilized or sterile sample inoculation on *A. adenophora* germination time (a, d), germination rate (b, e) and seedling survival (c, f) with phylogenetic distance from the inoculated local plant species to *A. adenophora*. Species effect (SE) = ln(conspecific/heterospecific) (details please see Methods). More negative SE values for germination time and positive SE values for germination rate and seedling survival represent more adverse effects of a local plant species on *A. adenophora*. The shaded areas around each line are 95% confidence intervals, and only significant linear relationships are shown (*P*<0.05). G0 and G21 represent plant tissue inoculation at G0 and G21 period, respectively. The phylogenetic distances from the inoculated local plant species to *A. adenophora* were calculated via the maximum composite likelihood model on the basis of Neighbour-joining (NJ) distance.

Correlation analysis revealed that leaf physical traits (DW, LA) and chemical traits (TP, TN) were related to aboveground bacterial communities, whereas leaf chemical traits (TP, TN) were related to aboveground fungal communities. Leaf physical traits (thickness) are related to aboveground fungal pathogen community. Specifically, leaf TN was negatively correlated with the germination rate (*P*<0.05). Leaf TP and TN were negatively correlated with seedling survival, but TA and TC were positively correlated with seedling survival (all *P*<0.05, Fig. 4). Aboveground microbial communities of plant species were significantly correlated with their effects on *A. adenophora* seedling survival (*P*<0.05) but not with germination (*P*>0.05); fungal (*P*<0.05), rather than bacterial pathogenic community (*P*>0.05), were correlated with seedling survival of *A. adenophora* (Fig. 4a, b). Additionally, local plants closer to *A. adenophora* showed more similar bacterial and fungal pathogen community composition to those in *A. adenophora* (Fig. 5a, b).

**Figure 4.**
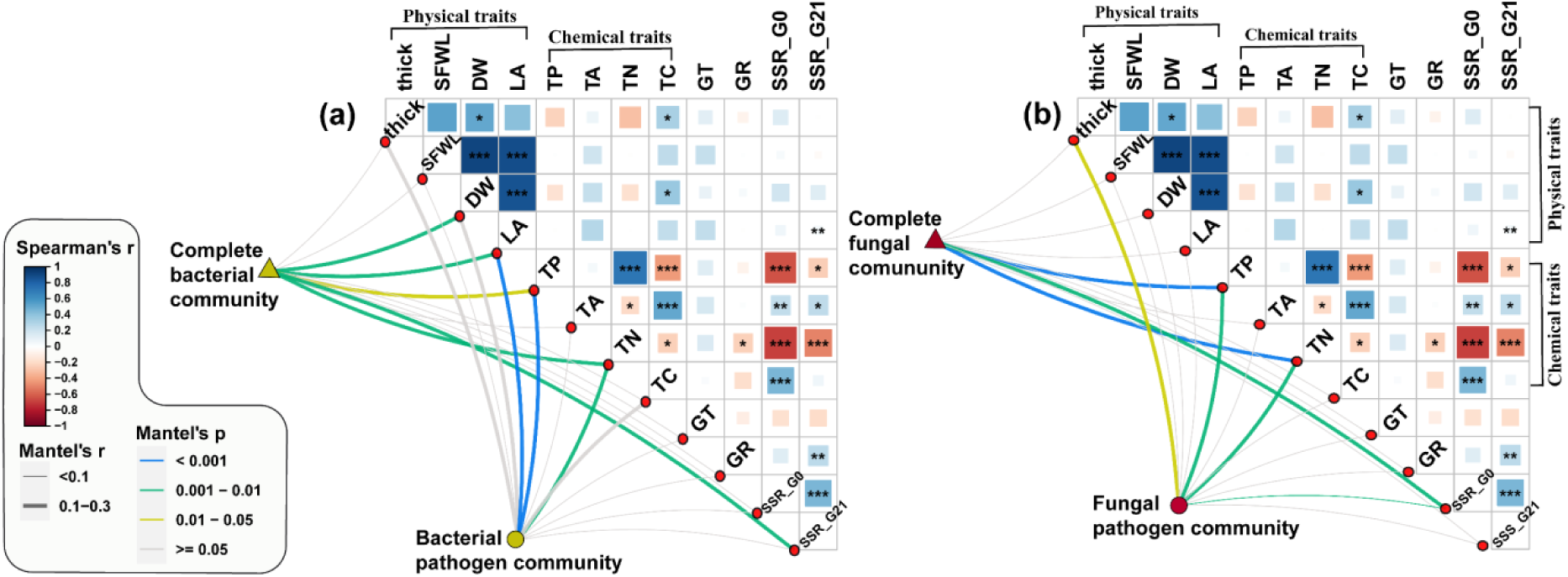
Relationships of bacterial (a) and fungal communities (b) with each leaf physiochemical trait, seed germination and seedling survival. The red and blue boxes represent negative and positive correlations, respectively. * *P* < 0.05, ** *P* < 0.01, *** *P* < 0.001. Edge width corresponds to Mantel’s r statistic, and edge colours denote statistical significance. Thick: leaf thickness, SFWL: saturated fresh weight of leaves, DW: leaf dry weight, LA: leaf area, TP: total phosphorus, TA: total polyphenols, TN: total nitrogen, TC: total carbon; GT and GR represent germination time, germination rate of unsterilized aboveground tissue inoculation, respectively; SSR_G0 and SSR_G21 represent seedling survival rate of unsterilized aboveground tissue inoculation at G0 and G21 period, respectively.

**Figure 5.**
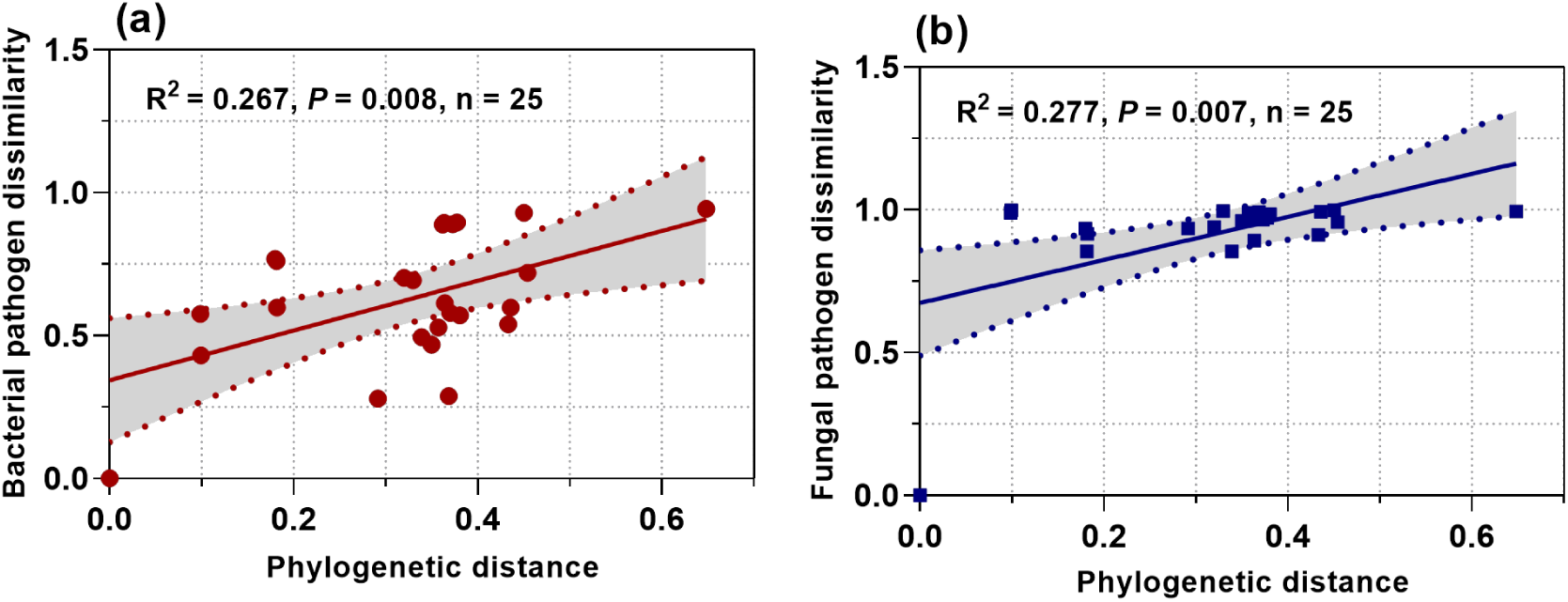
The relationship of plant phylogenetics with pathogen community dissimilarity in plant tissue. The relationship of phylogenetic distance from local plants to *A. adenophora* with bacterial (a) and fungal (b) pathogen community dissimilarity. The phylogenetic distances see Fig. 3. Bacterial and fungal pathogen community dissimilarity between local plant and *A. adenophora* were the Bray-Curtis distance calculated by average sequence numbers of bacterial and fungal pathogen ASVs in each plant species, respectively.

Aboveground microbial communities mainly explained seedling survival rather than germination (Fig. 4), thus, we further identify key microbes those correlated with seedling survival via random forest analysis. The 30 most predictive bacterial and fungal ASVs accounting for seedling survival were determined; all the bacterial ASVs (with the exception of BASV135_*Pseudomonas* and BASV80_*Caulobacter*) were positively related, but the fungal ASVs (with the exception of two Trichomeriaceae_unclassified ASVs and one *Phaeococcomyces*) were negatively related to seedling survival. Indeed, only two bacterial ASVs (BASV27_*Sphingomonas*, BASV57_*Pseudomonas*) and most fungal AVSs (e.g., members of *Didymella*, *Fusarium*, *Alternaria, Mycosphaerella, Stagonospora,* etc.) are plant pathogens. Interestingly, these fungal pathogen ASVs were mainly specialists (Fig. 6a-d).

**Figure 6.**
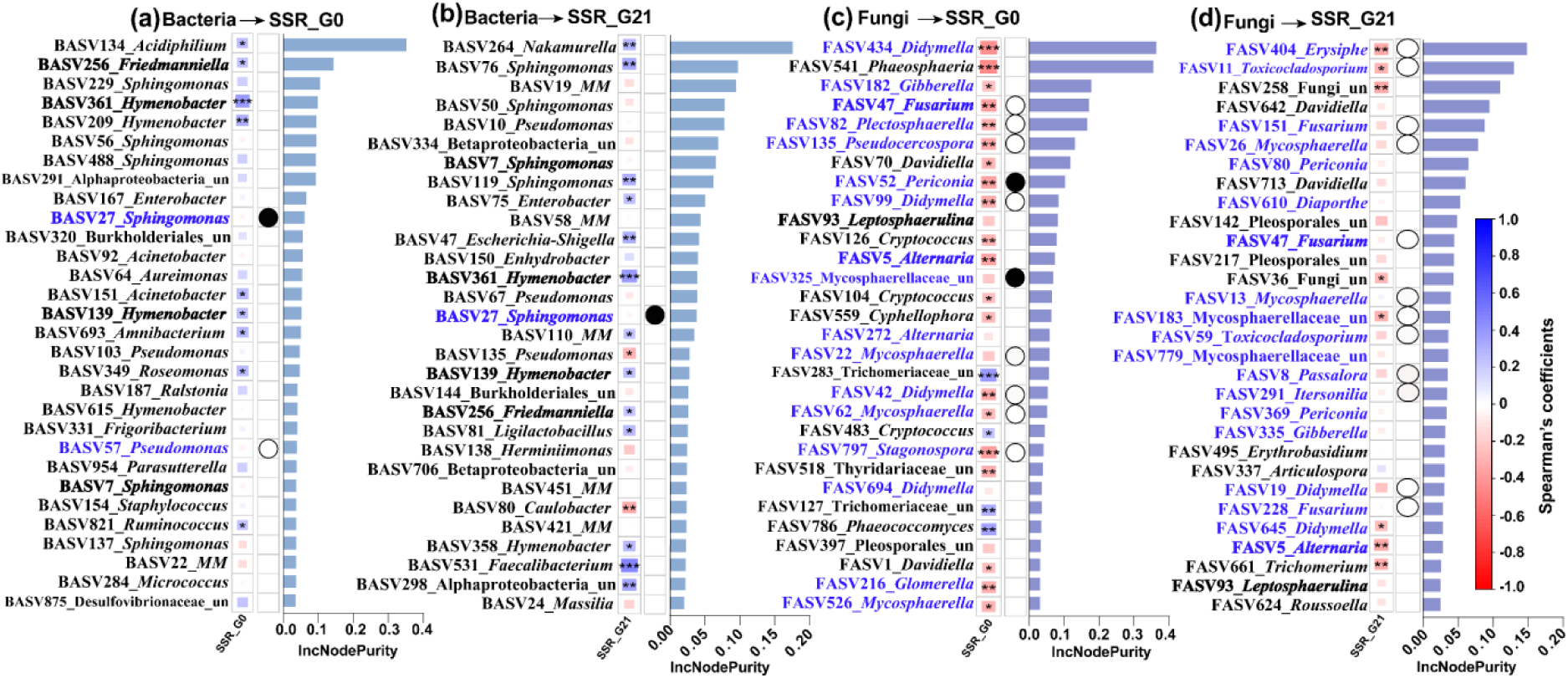
Random forest predictive bacterial and fungal ASVs accounting for seedling survival and their relationships with seedling survival. An increase in the node purity (IncNodePurity) of variables was used to estimate the importance of these predictors, where higher IncNodePurity values imply more important predictors. SSR_G0 and SSR_G21 represent microbial effects on seedling survival after plant tissue inoculation at G0 and G21 period, respectively. Microbial effect (ME) = ln(unsterilized/sterilized). The “un” in the figures is the abbreviation for “unclassified”, and “MM” is *Methylobacterium-Methylorubrum*. The blue font represents potential plant pathogen ASVs, and the bold font represents ASVs accounting for both SSR_G0 and SSR_G21. The solid and open circles represent generalist and specialist pathogen ASVs, respectively, and the non-mark is neither a generalist nor a specialist pathogen.

### 4. Identification of fungi related to seedling survival

We subsequently isolated 70 fungal strains from dead seedlings caused by aboveground tissue inoculation, and the numerically dominant genera were *Fusarium* (30 strains), *Chaetomium* (16 strains) and *Alternaria* (12 strains) (Fig. 7a, Table S4). The most (13 strains) fungi were isolated from dead seedlings inoculated with leaves of the plant species *Pueraria peduncularis*, followed by *Fallopia multiflora* (8 strains) and *Hypericum monogynum* (6 strains) (Fig. 7b). Seedling mortality caused by these fungi presented a significant phylogenetic signal (Pagel’s λ = 0.405, *P* < 0.0001). Overall, the highly virulent strains (> 60% mortality) belonged to the genera *Fusarium*, *Alternaria* and *Stagonosporopsis* (Fig. 7c).

**Figure 7.**
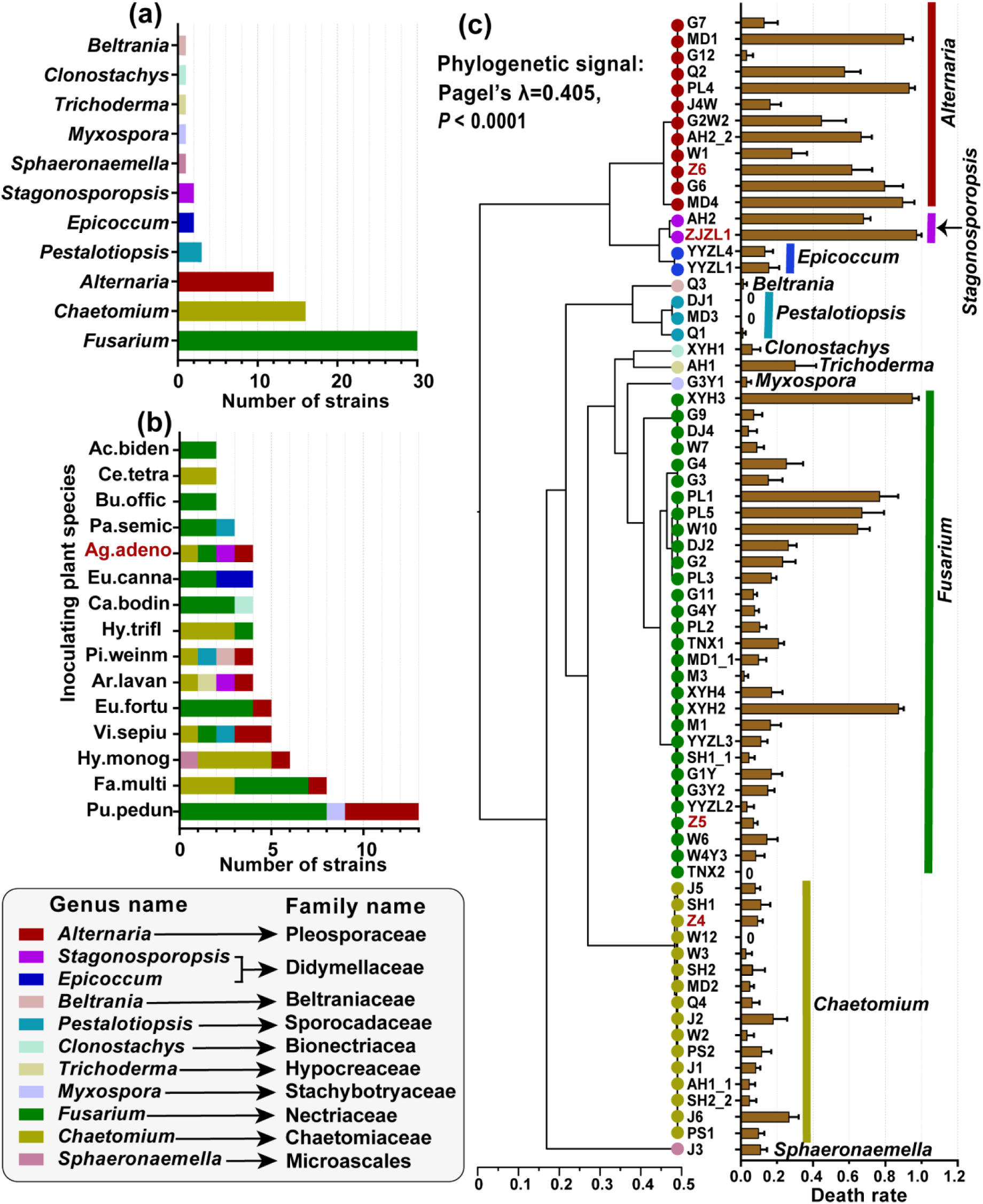
Cultivable fungi associated with dead *A. adenophora* seedlings inoculated with plant aboveground tissues on the 21^st^ day [G21] and their lethal effects on seedlings. Number of strains at the genus level (a). Number of strains isolated from dead seedlings inoculated with aboveground tissues of different plant species (b). The phylogenetic signals about the lethal effects of 70 fungal strains on *A. adenophora* seedlings (c). The red text highlights the exotic species *A. adenophora* and fungal strains isolated from dead seedlings inoculated with *A. adenophora* tissues. Means ± 1 SE, n = 5.

## Discussion

The resistance of local resident communities to plant invasion had not been characterized at the population establishment stage by comparing the roles of aboveground and belowground microbes. In this study, we compared the soil and aboveground tissue effects of 25 local plant species on the germination and survival of the invasive plant *A. adenophora*. We found that, compared with soil inoculation, aboveground tissue inoculation had more adverse effects on *A. adenophora* germination and seedling survival (Fig. 1). The greater relative abundance of plant pathogens in aboveground tissues than in soils could provide a reasonable explanation for this phenomenon (Fig. 2cd). Even so, it is also surprising that soils had relatively little effect on *A. adenophora* germination and seedling survival. Only sterile soil from one local plant species decreased the germination rate, and no species had a greater adverse effect on the germination time and seedling survival of *A. adenophora* when compared with control (i.e., physical and chemical effect), indirectly indicating that the soil allelopathy of local plants was negligible during the early growth stage of *A. adenophora*. Meanwhile, because unsterilized soil from all 26 plant species did not cause more significantly adverse effects on *A. adenophora* germination times, germination rates, and survival rates than their sterile soil (i.e., microbial effects) (Figs. 1 & S2), the edaphic microbial effects on *A. adenophora* germination and seedling survival were also negligible. Many studies on soil feedback have shown that invasive plant species commonly escape from soil pathogens in introduced regions (Callaway et al., 2011; Mitchell and Power, 2003). Here, it is unclear whether such negligible adverse effects on germination and seedlings in soil are due to release from soil-borne pathogens by *A. adenophora* because we have not tested the effects of the soil on local plant species themselves.

Further analysis revealed that leaf physicochemical traits and aboveground microbes of local plant species distinctively affected *A. adenophora* seed germination and seedling survival. Although most sterile aboveground tissue inoculations (physicochemical effects) of local plant species significantly delayed the germination time and decreased the germination rate than the control (Fig. S2), sterilization treatment did not significantly affect germination time or germination rate on the basis of generalized linear models (Table S2), and no unsterilized aboveground tissue caused more significantly adverse effects on *A. adenophora* germination time and germination rate than their sterile tissue (microbial effects) (Fig. S2). These findings suggest that leaf physicochemical traits, rather than aboveground microbes, have adverse effects on *A. adenophora* seed germination. In contrast, most unsterilized aboveground tissues but only a few sterile aboveground tissues resulted in significantly lower survival rates than did the control; a high proportion of plant species resulted in significantly lower seedling survival when comparing unsterilized aboveground tissue inoculations with sterile aboveground tissue inoculations (microbial effects) (Fig. 1). These data indicate that aboveground microbes rather than leaf physiochemical traits have adverse effects on seedling survival. Although aboveground microbiota effects on host health have been widely noted (Chen et al., 2020b; Vorholt, 2012), our study highlighted their important role in seedling survival.

Plant species trait similarity has been shown to be associated with phylogenetic relatedness (Fitzpatrick et al., 2017). Leaf physical and chemical traits and microbial communities varied among the 26 plant species in this study (Fig. S3), but plant phylogenetic distance did not account for the similarity of these leaf physical and chemical traits (Table S3). A possible reason may be that only a few plant traits were included in this study. Heavy natural selection may also be one possible reason for the inability to detect phylogenetic signals in the physicochemical traits of local plant species (Darwin, 1859). As in previous reports from tropical rainforests (Kembel and Mueller, 2014; Kembel et al., 2014), we also found that plant phylogenetic distance accounted for aboveground bacterial rather than fungal community dissimilarity (Table S3). Frequent fungal spore dispersal among host plants may contribute to the decreased phylogenetic relatedness of aboveground fungal community as a whole (Chen et al., 2020a). Interestingly, leaf chemical dissimilarity is positively correlated with aboveground fungal community dissimilarity, but physical dissimilarity is positively correlated with aboveground bacterial community dissimilarity (Table S3), suggesting that leaf functional traits can effectively shape the aboveground microbial community.

We found that only the germination rate was negatively correlated with phylogenetic distance caused by sterile inoculation (i.e., physical and chemical effects) (Fig. 3), and correlation analysis indicated that only total leaf nitrogen was negatively related to the germination rate. Previous studies have shown that relatively closely related plant leaves and shoots have relatively similar allelopathic potential and thus may have more adverse effects on closer plants (Grutters et al., 2017); high nutrient levels support strong allelochemical effects on seed germination (Wu et al., 2022). These data suggest that phylogenetic relatedness between local plants and the invasive *A. adenophora* mainly drives detrimental effects on the germination rate via similar allelopathy, which is supported mainly by the leaf N content.

With respect to seedling survival, the aboveground tissues of the native plants more closely related to those of *A. adenophora* more strongly decreased *A. adenophora* seedling survival in response to unsterilized inoculation (Fig. 3). Further correlation analysis revealed that the total aboveground microbial community, particularly the fungal pathogen community, was highly related to seedling survival, and leaf total nitrogen and phosphate were also negatively related to seedling survival (Fig. 4). Moreover, native plants more closely related to *A. adenophora* present more similar pathogenic community to those in *A. adenophora* (Fig. 5). These data suggest that phylogenetic relatedness between local plants and invaders has detrimental effects on seedling survival via aboveground fungal pathogens, which is supported mainly by high leaf N and P contents. These indicated that aboveground plant tissues with more similar nutrient contents share more pathogens. Similarly, closely related plant species often share more pathogens than distantly related species do (Burns and Strauss, 2011; Fitzpatrick et al., 2017; Gilbert and Webb, 2007), which is why phylogenetically distinct species in a community are more likely to invade successfully due to the lower susceptibility of plant pathogens (Parker et al., 2015). Plant phylogenetic relatedness commonly shapes soil-borne pathogen-driven seedling mortality (Huang et al., 2022; Packer and Clay, 2000). This is the first study to provide evidence for the effects of phylogenetic relatedness and leaf traits (both functional and microbial) on the local community resistance to *A. adenophora* invasion through the limitations of germination and seedling survival due to sharing similar aboveground fungal pathogen community.

We found that aboveground fungi, rather than bacteria, adversely impacted seedling survival (Figs. 4, 6, 7); unexpectedly, most of the detrimental fungal ASVs were more specialized (e.g., *Fusarium*, *Didymella, Mycosphaerella*) in the host range. Previous reports have indicated that invasive plants escape from specialist pathogens when they establish within the introduced range (Halbritter et al., 2012; Keane and Crawley, 2002). Our data indicated that, at the early seedling stage, *A. adenophora* did not escape from aboveground fungi associated with local plant species that have a limited host range (i.e., they are more specialized and only occur on a minority of plant species). Interestingly, many pathogen ASVs belonging to *Fusarium*, *Alternaria*, and *Stagonosporopsis* adversely impacted seedling survival (Figs. 6-7). These members have commonly been reported as generalist plant leafspot pathogens (Li et al., 2017; Omar et al., 2018; Vergnes et al., 2006); for example, *Alternaria* sp. is pathogenic to a large variety of plants, such as those causing stem cancer, leaf blight or leaf spot (Leiminger et al., 2015; Thomma, 2003; Vergnes et al., 2006). The highly lethal effects of these leafspot pathogens on young seedlings of *A. adenophora* in this case, on the one hand, suggest a previously unexpected role of these fungal species in seedling mortality of invader; on the other hand, these ASVs with high seedling-killing effects might actually represent some unknown new genotypes of these common genera. For example, one detrimental fungus, i.e., FASV5_*Alternaria*, occurs on all host plants, with high abundances on Asteraceae plants phylogenetically close to *A. adenophora* (e.g., *Eupatorium fortune* and *Eupatorium lindleyanum*) (See Fig. S4). It is valuable to verify whether FASV5_*Alternaria* is a potential novel species or genotype and whether the invader can benefit from seedling establishment via distinct lethal effects on invaders from local congeners elicited by this pathogen. Nonetheless, our method of using water agar plates to test seedling survival may overestimate their lethal effects. Indeed, many species or genotypes of *Fusarium* and *Alternaria* are also saprophytic (Summerell et al., 2011; Thomma, 2003). These fungi may act as pathogens in agar-grown seedlings.

The effects of plant phylogenetic relationships and leaf traits (both functional and microbial) on *A. adenophora* seedling survival were ontogeny dependent because plant phylogenetic distance was significantly negatively correlated with seedling survival when inoculating on day zero (SSR_G0) but not on day twenty-one (SSR_G21) (Fig. 3). Moreover, unsterilized plant tissues of most species resulted in lower seedling survival when inoculating on day zero than on day twenty-one (Fig. 1). Microbial impacts on their host are usually strongest in the early growth stages (Bagchi et al., 2014; Jevon et al., 2020). One potential reason is that small seedlings usually allocate most of their resources to survival and growth, whereas older seedlings have relatively more resources to defend against microbial infection (Geisen et al., 2021; Schloter and Matyssek, 2009). Another possible reason is related to beneficial effects of seed-borne bacteria on adverse external microbial sources in young seedlings. For example, seed endophytes could improve seedling resistance to abiotic and biotic stresses (Molina-Montenegro et al., 2023; Wang and Zhang, 2023). These seed endophytes might be inhibited or excluded from young seedlings by external sources of microbes inoculated on day zero, when seedlings are highly sensitive to external microbial infection (Geisen et al., 2021; Jevon et al., 2020). Therefore, determining how peripheral microbial sources interact with seed–borne endophytes across seedling ontogeny would be a valuable extension of our results. Taken together, our data suggest that the role of the phylogenetic relationships of local communities in *A. adenophora* should be more important for invasion resistance at earlier stages of invasion.

## Materials and methods

### Plant species and sample collection

Aboveground tissues and belowground soils from the invasive alien plant *Ageratina adenophora* and 25 native plants (belonging to 19 families) were collected in Kunming, Yunnan Province, China, in April 2022 (Table S1). For tree species, aboveground tissue and rhizosphere soil sample was collected from one randomly selected individual from one plot. We collected approximately 45 g of standing mature leaf (regardless of green or senescent, healthy or diseased spot leaves) rather than those fallen on the soil surface to minimize soil microbial contamination. For several forb and herb species (e.g. *Arthraxon prionodes*, *Vicia sepium*, *Senecio scandens*, and *Eupatorium lindleyanum*), due to their small biomass size we had to collect the aboveground tissues (including leaves, twigs, or possible inflorescences), following the methods previously described by Whitaker et al. (2017), from all individuals in a population from one plot. In total, five biologically independent aboveground tissues or belowground rhizosphere soils for each plant species were sampled from five plots (more than 100 m apart from each other). Rhizosphere soil (defined as that tightly attached to the roots) were collected by shaking roots. All the collected aboveground tissues and rhizosphere soils were air-dried in a clean room.

The soils were ground through a 2 mm sieve. Plant tissues were crushed into ∼2 mm fragments by fingers wearing sterile gloves. For those species with hard leaves, we had to cut continuously them into small fragments via sterile scissors. For convenience in the inoculation application, we packaged these samples in centrifuge tubes, each containing 0.1 g of plant tissue or soil, two copies per sample, where one copy was sterilized by gamma irradiation (30 kGy, 30 h, Huayuan Nuclear Radiation Technology Co., Ltd., Kunming, China) and one copy remained unsterilized, so that we could compare the microbial effects of the soil and aboveground tissues on the seed germination and seedling survival of *A. adenophora*. These tubes were finally stored at 4°C until inoculation. Another 0.3 g of plant tissues or soils from each sample were put into cryopreservation tubes at -80°C until DNA extraction for identifying microbial community. The remaining plant tissue was ground to a 0.15 mm sieve by a crusher for chemical content detection.

### Leaf functional trait measurements

For each plant species, we selected five fully developed, green and healthy leaves to measure the physical traits, such as leaf thickness (thick), leaf area (LA), saturated fresh weight of leaves (SFWL), and leaf dry weight (DW). We also measured leaf chemical traits, including total carbon (TC), total nitrogen (TN), total phosphorus (TP) and total polyphenols. Leaf TC and TN were measured via an elemental analyser (vario MACRO cube, Elementar, Germany); TP was measured via the molybdenum antimony colorimetric method. The total polyphenols were expressed as tannic acid equivalents (TAs), which were determined following the Folin-Ciocalteu assay referring to Ainsworth and Gillespie (2007).

### Inoculation experiment

Seeds of *A. adenophora* were collected from Xishan Forest Park, Kunming. Seeds were disinfected by dipping in 75% ethanol for 30 s, 2% NaOCl for 3 min, and then washing five times with sterile water before sowing. To quantify the effects of the aboveground tissue and soil microbes of *A. adenophora* or different local native plant species on *A. adenophora* seed germination and seedling survival, we placed 16 surface-sterilized *A. adenophora* seeds on a water agar (WA) plate and inoculated them with living or sterilized aboveground tissue or rhizosphere soil across 26 plant species (hereafter referred to as G0 inoculation). We also included a plant tissue-free and soil-free control group to observe possible physical and/or chemical effects compared with sterilized samples. This resulted in a total of 525 plates (species [26] * soil vs. leaf [2] * sterile vs. unsterilized [2] * replicate [5] + control [5]). When inoculated, 0.1 g of plant tissues or rhizosphere soils (ground to 2 mm) were evenly distributed to directly touch each of the 16 seeds in a WA plate (see Figure S1). All plates were randomly placed in growth chambers (RXZ-380D, Ningbo Southeast Instrument Co., Ltd., Ningbo, China) and rearranged every week to mitigate potential positional effects. The chambers were kept at a temperature of 25/20°C (day/night), a light intensity of 12 000 lux (12 h/12 h) and a humidity of 65%. The number of germinated seeds was recorded every day for the first 14 days to calculate the germination time (GT), and the total number of germinated seeds on the 21^st^ day was recorded to calculate the final germination rate (GR). GT was calculated via the formula GT= Σ(G*i*×*i*)/ΣG*i* (*i*: number of days between seed sowing (day 0) and seed germination; G*i*: number of seeds germinated on day *i*) (Zhang et al., 2014). The GR was calculated as the proportion of seeds that had germinated by the 21^st^ day. The number of surviving seedlings on the 28^th^ day was used to calculate the seedling survival rate of the G0 inoculation (named SSR_G0). Because we observed high seedling mortality with unsterilized plant tissue inoculation, we performed an additional inoculation on 21-day-old control seedlings to explore the resistance of *A. adenophora* to aboveground microbes across seedling ontogeny (hereafter referred to as G21 inoculation), resulting in a total of 265 plates (species [26] * sterile vs. unsterilized [2] * replicate [5] + control [5]). We used the proportion of seedlings alive 7 days later to calculate the seedling survival rate (hereafter referred to as SSR_G21).

### Detection of microbial community in the inoculation sources

Aboveground plant tissue and belowground soil bacterial and fungal communities associated with 26 plant species were identified via Illumina sequencing, and each species including three independently biological duplications were sequenced. For DNA extraction, target‒gene amplification and sequencing details, see Supplementary Methods 1. The aboveground bacterial community of *Pistacia weinmannifolia* and the belowground fungal community of *Eupatorium lindleyanum* were not successfully sequenced three times, so analyses of the microbial communities did not include them.

### Plant phylogeny

Three genes (two plastid genes and one nuclear gene) for each species were obtained from GenBank or from sequencing (those not found in GenBank): ribulose-bisphosphate carboxylase (*rbc*L), maturase K (*mat*K), and the internal transcribed spacer (ITS) adjacent to the 5.8S ribosomal RNA gene (Table S1). The sequences were aligned in MEGA via MUSCLE with default parameters (Edgar, 2004; Tamura et al., 2013), followed by manual cutting and alignment checking. The aligned sequences were concatenated (*rbcL*–*matK*–ITS) via BioEdit (Hall, 1999). We calculated the pairwise phylogenetic distances on the basis of Neighbour-Joining (NJ) distance using Maximum Composite Likelihood model among all pairs of 26 plant species via MEGA.

### Seedling-killing fungus experiment

Although unsterilized plant tissue inoculation on day zero (G0 experiment) caused high seedling mortality (see Fig. 1, S1), dead seedlings had decomposed by the end of that experiment, so we isolated fungi from dead seedlings 7 d after inoculated 0.1 g of plant tissue fragments with 21-day-age seedlings (G21 experiment). In brief, each dead seedling was cut into approximately 1 mm × 1 mm pieces, and three pieces were placed into a Potato Dextrose Agar (PDA) plate and incubated at 20–25°C (day/night, 12 h/12 h) for 6–8 days or until mycelia grew. The hyphal tips were subsequently transferred onto new PDA plates and incubated until pure colonies appeared. The DNA of these purified fungal strains was extracted and identified by sequencing the ITS region (for sequencing details, see Supplementary Method S1).

Furthermore, we tested the effects of these strains on *A. adenophora* seedling survival. Sixteen surface-sterilized *A. adenophora* seeds were sown in a water agar plate. All 21-day-old seedlings in one water agar plate were inoculated with one fungal strain by touching 3 mm diameter agar discs with fungal mycelia to the leaves and stems. Five plates were used as five replicates for each strain. Fungi were grown on PDA plates for 7 days in an incubator at 25°C in the dark before inoculation. Seedlings were considered dead when the leaf and stem became brown and rotted. We calculated the mortality rate as the proportion of seedlings that died in a plate 14 d after inoculation. To examine the phylogenetic signal of the seedling-killing effects of fungal strains on *A. adenophora*, we calculated Pagel’s λ with the R package ‘phytools’, which measures the distribution of a trait across a phylogeny. A Pagel’s λ closer to 1 and *P* < 0.05 indicated a stronger phylogenetic signal. BEAST v. 1.10.4 was used to construct a Bayesian phylogenetic tree via these fungal sequences (Drummond et al., 2012). The resulting tree was visualized in FigTree v.1.4.3.

### Statistical analysis

Generalized linear models (GLMs) with quasipoisson distributions (link: log) (for GR and SSR based on germinated number and survival number) or Gaussian distributions (link: identity) (for GT) were used to compare the effects of sample type (plant tissue vs. soil), plant species, sterilization treatments on GT, GR and SSR via the R 4.2.0 package “lme4” (Bates et al., 2015). *P* values were estimated via the ANOVA function via chi-square (χ^2^) tests in GLMs. Nonparametric Mann–Whitney U tests were used to identify differences between each species in the unsterilized or sterilized treatments and the control, as well as the differences between each species in the unsterilized and sterilized treatments. The Kruskal‒Wallis test was performed to compare the differences among the four treatment groups of one species. The nonparametric Mann–Whitney U test and Kruskal‒Wallis test were performed using SPSS v. 27.0.

Nonmetric multidimensional scaling (NMDS) analysis and permutational analysis of variance (PERMANOVA) with the ANOSIM function in the R package “vegan” were also conducted to test the differences in physical and chemical traits on the basis of Euclidean distance and in the bacterial and fungal communities and functions on the basis of Bray‒Curtis distance among the plant species. Bacterial functional profiles were predicted via functional annotation of prokaryotic taxa (FAPROTAX) (Louca et al., 2016). Fungal functional guilds were inferred via FUNGuild, and guild assignments with confidence rankings of “highly probable” and “probable” were retained (Nguyen et al., 2016). Sequences that had multiple functional assignments in FUNGuild were excluded from the analysis. Those ASVs assigned as “plant pathogen” via FAPROTAX or FUNGuild were used as pathogen ASVs for further focusing on bacterial and fungal pathogen community. We also calculated relative abundance of these ASVs belonging to plant pathogens in each plant tissue sample. Plant pathogen ASVs were further classified into specialist/generalist pathogens via the “EcolUtils” package in R version 4.3.3 on the basis of niche width and permutation algorithms (Tanunchai et al., 2022).

We assumed that the distinct effects of aboveground tissues from each local plant species on *A. adenophora* seed germination and seedling survival manifested as phylogenetic distance. Therefore, when to determine whether phylogenetic distance was correlated with *A. adenophora* seed germination and seedling survival, we calculated the local plant species effect (SE) from aboveground tissues as the ratio of seed germination or seedling survival when inoculated with the *A. adenophora* (conspecific) tissue samples versus the heterospecific (local plant) tissue samples on the basis of a previously described method (see Semchenko et al. (2018)). For the germination rate (GR) and time (GT): Species effect (SE) = ln(conspecific/heterospecific). Because some plates had no surviving seedlings, for seedling survival rate (SSR): Species effect (SE) = ln([# SSR in conspecific +1]/[# SSR in heterospecific +1]). More positive SE values for germination rate and seedling survival and more negative SE values for germination time represent more adverse effects of a local plant species on *A. adenophora*.

To define the relationship of local plant species effects from aboveground tissues on *A. adenophora* germination and survival with plant phylogenetics, physical and chemical traits and microbial community, the paired phylogenetic distance was neighbour-joining (NJ) distance via the maximum composite likelihood model on the basis of concatenated *rbcL*–*matK*–ITS sequences; the physical distance of all pairs of plant species was the calculated Euclidean distance on the basis of LA, thickness, SFWL and DW; and the chemical distance was the calculated Euclidean distance on the basis of TA, TC, TN and TP. For fungal and bacterial complete community dissimilarity, the Bray‒Curtis distance was calculated on the basis of fungal and bacterial ASVs sequence number in all samples. The Bray‒ Curtis distance was calculated on the basis of fungal and bacterial pathogen ASVs sequence numbers in all plant tissue samples to characterize fungal and bacterial pathogen community dissimilarity.

We correlated the paired phylogenetic distance from *A. adenophora* to each local plant species with the calculated species effects (SE) on the seed germination and seedling survival of invasive *A. adenophora* via linear regression. We subsequently used Mantel tests to determine the correlations between host plant phylogenetic distance and leaf functional traits (i.e., leaf physical and chemical traits and the leaf bacterial and fungal communities) via the “ade4” package. In addition, a heatmap was generated, and a Mantel test was performed for visualization via the R “linkET”, “dplyr”, and “ggplot2” packages to determine the relationships among each leaf trait, seed germination, and seedling survival with complete microbial communities and pathogen communities, in which the Bray-Curtis distance algorithm was used to conduct Mantel tests on complete and pathogenic microbial communities. We identified the key fungal and bacterial predictive ASVs for seedling survival in response to G0 and G21 inoculation by random forest analysis on the basis of the node purity (IncNodePurity) index via the “randomForest” package in R 4.2.0, higher IncNodePurity values imply more important predictors. We calculated the microbial effect (ME) of aboveground tissue on *A. adenophora* seedling survival as the ratio of seedling survival when inoculated with the unsterilized samples versus the sterilized samples. Microbial effect (ME) = ln([# SSR in unsterilized +1]/[# SSR in sterilized +1]). A ME value greater than or less than zero represents positive or negative effects of the microbial community associated with the inoculum plant species on *A. adenophora* seedling survival. Finally, we correlated the relative abundance of such predictive ASVs with microbial effect on *A. adenophora* seedling survival according to Spearman’s correlation analysis.

## Conclusion

We evaluated the effects of the aboveground and belowground microbes and foliar functional traits of 25 native plants on the invasive *Ageratina adenophora*. We found that aboveground tissues had more adverse effects on *A. adenophora* establishment by delaying germination time and decreasing the germination rate and seedling survival than did soils; aboveground tissues from native plants, which are more closely related to *A. adenophora*, cause longer germination times and greater seedling mortality of *A. adenophora*. Local plants more closely related to *A. adenophora* present more similar pathogenic community to those in *A. adenophora*. Moreover, local plant aboveground fungi, rather than bacteria, adversely impact seedling survival, and most of the detrimental fungal pathogen ASVs are specialists. Several strains isolated from dead seedlings caused by aboveground tissue inoculation belonging to *Alternaria*, *Fusarium* and *Stagonosporopsis* were verified to kill more than 60% of *A. adenophora* seedlings and thus have great potential as biocontrol agents for *A. adenophora*. Our results highlight the effects of native plant aboveground microbes on preventing alien establishment, especially those closely related to alien plants. This study provides valuable information and possibilities for controlling alien plant invasion through the assembly of local plant communities and plant aboveground microbes.

## Supporting information

The supplementary material includes one supplementary method, four figures and four tables.

## Acknowledgements

We thank Xiao-Han Jin, Yu Li, Jin-peng Li, and Lu Cheng for their help with sample collection and experimental performance. This work was supported by the National Key R&D Program of China (*grant numbers 2022YFC2601100*), the Major Science and Technology Project in Yunnan Province, PR China (*grant numbers 202301AS070023*), and the “Double First-Class” University Project of Yunnan University.

## Competing Interests

We declare that we have no competing interests.

## Author contributions

Z.Z., H.Z., B.L. designed the research. Z.Z., Z.L., A.Y., Y.L. and Y.W. performed experiments. Z.Z. performed data analyses. Z.Z., H.Z. wrote the paper. B.L. and E.S. revised the manuscript. All authors read and approved the contents of this paper.

## Data availability

The data that support the findings of this study are available in NCBI at GenBank under accession numbers PRJNA1034946, PRJNA1034808 or via link https://www.ncbi.nlm.nih.gov/bioproject/PRJNA1034946 and https://www.ncbi.nlm.nih.gov/bioproject/PRJNA1034808 for leaf litter samples, PRJNA1182656, PRJNA1185034 or via link https://www.ncbi.nlm.nih.gov/bioproject/PRJNA1182656 and https://www.ncbi.nlm.nih.gov/bioproject/PRJNA1185034 for soil samples, respectively. Fungal ITS sequences were obtained from GenBank under accession numbers OR762577-OR762646 or via link https://www.ncbi.nlm.nih.gov/nuccore/?term=OR762577:OR762646[accn]. The other data and all analysis code deposited in the Dryad Digital Repository: https://datadryad.org/stash/share/Y8Xf141GOIEtK5tFa3nUZbyIOsagQDXzpfsT2ShPgh0.

## Supplementary Information

The supplementary material includes one supplementary method, four figures and four tables.

### Supplementary methods

**Method S1:** DNA extraction, target‒gene amplification and sequencing.

### Supplementary figures

**Figure S1.** Seedlings were inoculated with soils or plant tissues (from four randomly selected species) 21 days after sowing.

**Figure S2.** Effects of sterile and unsterilized plant tissue or soil inoculation on *A. adenophora* germination time (a) and germination rate (b) among the 26 species.

**Figure S3.** Leaf physical (a) and chemical (b) trait dissimilarity for 26 plant species.

**Figure S4.** The relative abundance of predictive fungal ASVs significantly correlated with seedling survival associated with aboveground tissues of 26 species.

### Supplementary tables

**Table S1.** Information for sample collection.

**Table S2.** Effects of plant sample type, species and sterilization on the germination and survival of *A. adenophora*.

**Table S3.** Mantel test results showing the relationships between host phylogenetic, physical or chemical distance and plant tissue microbial community dissimilarity.

**Table S4.** Detailed information on 70 fungal strains isolated from dead seedlings.

## Notes

https://www.ncbi.nlm.nih.gov/bioproject/PRJNA1034946

https://www.ncbi.nlm.nih.gov/bioproject/PRJNA1034808

https://www.ncbi.nlm.nih.gov/bioproject/PRJNA1182656

https://www.ncbi.nlm.nih.gov/bioproject/PRJNA1185034

### Competing Interest Statement

The authors have declared no competing interest.

### Summary of Updates

We added some detail for some Method part, and we creat a new figure (i.e. Figure 5) in this submission.

## References

Ainsworth, E.A., and Gillespie, K.M. (2007). Estimation of total phenolic content and other oxidation substrates in plant tissues using Folin–Ciocalteu reagent. Nature Protocols 2: 875–877.

Bagchi, R., Gallery, R.E., Gripenberg, S., Gurr, S.J., Narayan, L., Addis, C.E., Freckleton, R.P., and Lewis, O.T. (2014). Pathogens and insect herbivores drive rainforest plant diversity and composition. Nature 506: 85-+.

Bates, D., Machler, M., Bolker, B.M., and Walker, S.C. (2015). Fitting Linear Mixed-Effects Models Using lme4. Journal of Statistical Software 67: 1–48.

Burns, J.H., and Strauss, S.Y. (2011). More closely related species are more ecologically similar in an experimental test. Proc Natl Acad Sci U S A 108: 5302–5307.

Callaway, R.M., Bedmar, E.J., Reinhart, K.O., Silvan, C.G., and Klironomos, J. (2011). Effects of soil biota from different ranges on Robinia invasion: acquiring mutualists and escaping pathogens. Ecology 92: 1027–1035.

Chen, L., Zhou, J., Zeng, T., Miao, Y.-F., Mei, L., Yao, G.-B., Fang, K., Dong, X.-F., Sha, T., Yang, M.-Z., Li, T., Zhao, Z.-W., and Zhang, H.-B. (2020a). Quantifying the sharing of foliar fungal pathogens by the invasive plant *Ageratina adenophora* and its neighbours. New Phytologist 227: 1493–1504.

Chen, T., Nomura, K., Wang, X.L., Sohrabi, R., Xu, J., Yao, L.Y., Paasch, B.C., Ma, L., Kremer, J., Cheng, Y.T., Zhang, L., Wang, N.A., Wang, E.T., Xin, X.F., and He, S.Y. (2020b). A plant genetic network for preventing dysbiosis in the phyllosphere. Nature 580: 653-+.

Darwin, C. (1859). On the Origin of the Species by Means of Natural Selection: or,the Preservation of Favoured Races in the Struggle for Life. London: John Murray.

Demey, A., Staelens, J., Baeten, L., Boeckx, P., Hermy, M., Kattge, J., and Verheyen, K. (2013). Nutrient input from hemiparasitic litter favors plant species with a fast-growth strategy. Plant and Soil 371: 53–66.

Drummond, A.J., Suchard, M.A., Xie, D., and Rambaut, A. (2012). Bayesian Phylogenetics with BEAUti and the BEAST 1.7. Molecular Biology and Evolution 29: 1969–1973.

Edgar, R.C. (2004). MUSCLE: multiple sequence alignment with high accuracy and high throughput. Nucleic Acids Res. 32: 1792–1797.

Emmett, B.D., Youngblut, N.D., Buckley, D.H., and Drinkwater, L.E. (2017). Plant phylogeny and life history shape rhizosphere bacterial microbiome of summer annuals in an agricultural field. Frontiers in microbiology 8.

Fang, K., Chen, L., Zhou, J., Yang, Z.P., Dong, X.F., and Zhang, H.B. (2019). Plant-soil-foliage feedbacks on seed germination and seedling growth of the invasive plant *Ageratina adenophora*. Proceedings of the Royal Society B-Biological Sciences 286.

Fitzpatrick, C.R., Gehant, L., Kotanen, P.M., and Johnson, M.T.J. (2017). Phylogenetic relatedness, phenotypic similarity and plant-soil feedbacks. Journal of Ecology 105: 786–800.

Geisen, S., ten Hooven, F.C., Kostenko, O., Snoek, L.B., and van der Putten, W.H. (2021). Fungal root endophytes influence plants in a species-specific manner that depends on plant’s growth stage. JOURNAL OF ECOLOGY 109: 1618–1632.

Gilbert, G.S., Magarey, R., Suiter, K., and Webb, C.O. (2012). Evolutionary tools for phytosanitary risk analysis: phylogenetic signal as a predictor of host range of plant pests and pathogens. Evolutionary Applications 5: 869–878.

Gilbert, G.S., and Webb, C.O. (2007). Phylogenetic signal in plant pathogen-host range. Proceedings of the National Academy of Sciences of the United States of America 104: 4979–4983.

Grutters, B.M.C., Saccomanno, B., Gross, E.M., Van de Waal, D.B., van Donk, E., and Bakker, E.S. (2017). Growth strategy, phylogeny and stoichiometry determine the allelopathic potential of native and non-native plants. Oikos 126: 1770–1779.

Gu, C., Tu, Y., Liu, L., Wei, B., Zhang, Y., Yu, H., Wang, X., Yangjin, Z., Zhang, B., and Cui, B. (2021). Predicting the potential global distribution of Ageratina adenophora under current and future climate change scenarios. Ecology and Evolution 11: 12092–12113.

Halbritter, A.H., Carroll, G.C., Güsewell, S., and Roy, B.A. (2012). Testing assumptions of the enemy release hypothesis: generalist versus specialist enemies of the grass *Brachypodium sylvaticum*. Mycologia 104: 34–44.

Hall, T. (1999). BioEdit: a user-friendly biological sequence alignment editor and analysis program for Windows 95/98/NT. Nucleic Acids Symposium Series 41: 95-98.

He, Y.F., Jia, B.B., Wei, C.Q., Fan, F.Y., Wilschut, R.A., and Lu, X.M. (2023). Leaf litter presence in the non-growing season prolongs plant legacy effects on soil fungal communities and succeeding plant growth. Journal of Ecology 111: 1997–2009.

Huang, T.H., Huang, C.L., Lin, Y.C., and Sun, I.F. (2022). Seedling survival simultaneously determined by conspecific, heterospecific, and phylogenetically related neighbors and habitat heterogeneity in a subtropical forest in Taiwan. Ecology and Evolution 12.

Huang, Z.Y., Liu, S.S., Bradford, K.J., Huxman, T.E., and Venable, D.L. (2016). The contribution of germination functional traits to population dynamics of a desert plant community. Ecology 97: 250–261.

Jessen, M.-T., Auge, H., Harpole, W.S., and Eskelinen, A. (2023). Litter accumulation, not light limitation, drives early plant recruitment. Journal of Ecology 111: 1174–1187.

Jevon, F.V., Record, S., Grady, J., Lang, A.K., Orwig, D.A., Ayres, M.P., and Matthes, J.H. (2020). Seedling survival declines with increasing conspecific density in a common temperate tree. Ecosphere 11: e03292.

Keane, R.M., and Crawley, M.J. (2002). Exotic plant invasions and the enemy release hypothesis. Trends in Ecology & Evolution 17: 164–170.

Kembel, S.W., and Mueller, R.C. (2014). Plant traits and taxonomy drive host associations in tropical phyllosphere fungal communities. Botany 92.

Kembel, S.W., O’Connor, T.K., Arnold, H.K., Hubbell, S.P., Wright, S.J., and Green, J.L. (2014). Relationships between phyllosphere bacterial communities and plant functional traits in a neotropical forest. Proceedings of the National Academy of Sciences of the United States of America 111: 13715–13720.

Kimball, S., Angert, A.L., Huxman, T.E., and Venable, D.L. (2010). Contemporary climate change in the Sonoran Desert favors cold-adapted species. Global Change Biology 16: 1555–1565.

Lamb, E.G. (2008). Direct and indirect control of grassland community structure by litter, resources, and biomass. Ecology 89: 216–225.

Leiminger, J., Bassler, E., Knappe, C., Bahnweg, G., and Hausladen, H. (2015). Quantification of disease progression of Alternaria spp. on potato using real-time PCR. European Journal of Plant Pathology 141: 295–309.

Li, H.-X., Gottilla, T.M., and Brewer, M.T. (2017). Organization and evolution of mating-type genes in three *Stagonosporopsis* species causing gummy stem blight of cucurbits and leaf spot and dry rot of papaya. Fungal Biology 121: 849–857.

Li, S.P., Cadotte, M.W., Meiners, S.J., Hua, Z.S., Shu, H.Y., Li, J.T., and Shu, W.S. (2015). The effects of phylogenetic relatedness on invasion success and impact: deconstructing Darwin’s naturalisation conundrum. Ecology Letters 18: 1285–1292.

Liu, X.B., Liang, M.X., Etienne, R.S., Wang, Y.F., Staehelin, C., and Yu, S.X. (2012). Experimental evidence for a phylogenetic Janzen-Connell effect in a subtropical forest. Ecology Letters 15: 111–118.

Louca, S., Parfrey, L.W., and Doebeli, M. (2016). Decoupling function and taxonomy in the global ocean microbiome. SCIENCE 353: 1272–1277.

Márquez, S.S., Bills, G.F., and Zabalgogeazcoa, I. (2011). Fungal species diversity in juvenile and adult leaves of *Eucalyptus globulus* from plantations affected by Mycosphaerella leaf disease. Annals of Applied Biology 158: 177–187.

Mitchell, C.E., and Power, A.G. (2003). Release of invasive plants from fungal and viral pathogens. Nature 421: 625–627.

Möhler, H., Diekötter, T., Herrmann, J.D., and Donath, T.W. (2018). Allelopathic vs. autotoxic potential of a grassland weed—evidence from a seed germination experiment. Plant Ecology & Diversity 11: 539–549.

Molina-Montenegro, M.A., Ballesteros, G.I., Acuña-Rodríguez, I.S., Pertierra, L.R., Greve, M., Richardson, D.M., Convey, P., Biersma, E.M., Goodall-Copestake, W.P., and Newsham, K.K. (2023). The "Trojan horse" strategy: Seed fungal endophyte symbiosis helps to explain the invasion success of the grass, *Poa annua*, in Maritime Antarctica. Diversity and Distributions.

Nguyen, N.H., Song, Z.W., Bates, S.T., Branco, S., Tedersoo, L., Menke, J., Schilling, J.S., and Kennedy, P.G. (2016). FUNGuild: An open annotation tool for parsing fungal community datasets by ecological guild. FUNGAL ECOLOGY 20: 241–248.

Omar, N.H., Mohd, M., Mohamed Nor, N.M.I., and Zakaria, L. (2018). Characterization and pathogenicity of *Fusarium* species associated with leaf spot of mango (*Mangifera indica* L.). Microbial Pathogenesis 114: 362–368.

Packer, A., and Clay, K. (2000). Soil pathogens and spatial patterns of seedling mortality in a temperate tree. Nature 404: 278–281.

Parker, I.M., Saunders, M., Bontrager, M., Weitz, A.P., Hendricks, R., Magarey, R., Suiter, K., and Gilbert, G.S. (2015). Phylogenetic structure and host abundance drive disease pressure in communities. Nature 520: 542–544.

Schloter, M., and Matyssek, R. (2009). Tuning growth versus defence–belowground interactions and plant resource allocation. Plant and Soil 323: 1–5.

Semchenko, M., Leff, J.W., Lozano, Y.M., Saar, S., Davison, J., Wilkinson, A., Jackson, B.G., Pritchard, W.J., De Long, J.R., Oakley, S., Mason, K.E., Ostle, N.J., Baggs, E.M., Johnson, D., Fierer, N., and Bardgett, R.D. (2018). Fungal diversity regulates plant-soil feedbacks in temperate grassland. Sci Adv 4: eaau4578.

Summerell, B.A., Leslie, J.F., Liew, E.C.Y., Laurence, M.H., Bullock, S., Petrovic, T., Bentley, A.R., Howard, C.G., Peterson, S.A., Walsh, J.L., and Burgess, L.W. (2011). *Fusarium* species associated with plants in Australia. Fungal Diversity 46: 1–27.

Tamura, K., Stecher, G., Peterson, D., Filipski, A., and Kumar, S. (2013). MEGA6: Molecular Evolutionary Genetics Analysis Version 6.0. Molecular Biology and Evolution 30: 2725–2729.

Tanunchai, B., Ji, L., Schroeter, S.A., Wahdan, S.F.M., Larpkern, P., Lehnert, A.S., Alves, E.G., Gleixner, G., Schulze, E.D., Noll, M., Buscot, F., and Purahong, W. (2022). A poisoned apple: First insights into community assembly and networks of the fungal pathobiome of healthy-looking senescing leaves of temperate trees in mixed forest ecosystem. Frontiers in Plant Science 13.

Thomma, B.P. (2003). Alternaria spp.: from general saprophyte to specific parasite. Molecular plant pathology 4: 225–236.

van Kleunen, M., Bossdorf, O., and Dawson, W. (2018). The ecology and evolution of alien plants. In: Annual Review of Ecology, Evolution, and Systematics, Vol 49--Futuyma, D.J., ed. 25-47.

Vergnes, D.M., Renard, M.E., Duveiller, E., and Maraite, H. (2006). Identification of Alternaria spp. on wheat by pathogenicity assays and sequencing. Plant Pathology 55: 485–493.

Vorholt, J.A. (2012). Microbial life in the phyllosphere. Nature Reviews Microbiology 10: 828–840.

Wang, R., and Wang, Y.Z. (2006). Invasion dynamics and potential spread of the invasive alien plant species Ageratina adenophora (Asteraceae) in China. Diversity and Distributions 12: 397–408.

Wang, Y.-L., and Zhang, H.-B. (2023). Assembly and function of seed endophytes in response to environmental stress. Journal of microbiology and biotechnology 33: 1–11.

Whitaker, B.K., Bauer, J.T., Bever, J.D., and Clay, K. (2017). Negative plant-phyllosphere feedbacks in native Asteraceae hosts - a novel extension of the plant-soil feedback framework. Ecol Lett 20: 1064–1073.

Wu, C.Y., Chen, Y.F., and Grenier-Héon, D. (2022). Effects of allelopathy and availability of nutrients and water resources on the survival and growth of plant species in a natural Dacrydium forest. Canadian Journal of Forest Research 52: 100–108.

Xiong, S., and Nilsson, C. (1999). The effects of plant litter on vegetation: a meta-analysis. Journal of Ecology 87: 984–994.

Zeng, Z.-Y., Huang, J.-R., Liu, Z.-Q., Yang, A.-L., Li, Y.-X., Wang, Y.-L., and Zhang, H.-B. (2024). Distinct effects of phyllosphere and rhizosphere microbes on invader *Ageratina adenophora* during its early life stages. eLife: 2024.2001.2009.574925.

Zeng, Z.Y., Yang, Z.P., Yang, A.L., Li, Y.X., and Zhang, H.B. (2023). Genetic evidence for *Colletotrichum gloeosporioides* transmission between the invasive plant *Ageratina adenophora* and co-occurring neighbor plants. Microbial Ecology 86: 2192–2201.

Zhang, C., Willis, C.G., Burghardt, L.T., Qi, W., Liu, K., de Moura Souza-Filho, P.R., Ma, Z., and Du, G. (2014). The community-level effect of light on germination timing in relation to seed mass: a source of regeneration niche differentiation. New Phytol 204: 496–506.

Zhang, R., Hu, X., Baskin, J.M., Baskin, C.C., and Wang, Y. (2017). Effects of Litter on Seedling Emergence and Seed Persistence of Three Common Species on the Loess Plateau in Northwestern China. Frontiers in plant science 8: 103.

Zheng, Y.L., Burns, J.H., Liao, Z.Y., Li, Y.P., Yang, J., Chen, Y.J., Zhang, J.L., and Zheng, Y.G. (2018). Species composition, functional and phylogenetic distances correlate with success of invasive *Chromolaena odorata* in an experimental test. Ecology Letters 21: 1211–1220.

