## Supplementary material for "Aboveground fungal lethal effects of local plants on invasive plant *Ageratina adenophora* seedlings": The supplementary material includes one supplementary method, four figures and four tables.: Supplementary for 20250412.pdf

### Supplementary information

### 1. Supplementary methods

#### Method S1: DNA extraction, target–gene amplification and sequencing

Total DNA was extracted from the plant tissue using the cetyltrimethylammonium bromide (CTAB) method (Stewart and Via, 1993). The quality of the extracted DNA was assessed by electrophoresis in a 1.5% agarose gel using an ND-1000 spectrophotometer (NanoDrop Technology, Wilmington, USA). A Qubit dsDNA HS assay kit (Invitrogen, USA) was used to quantify the DNA concentration. We amplified the bacterial 16S rRNA V4 region and fungal ITS2 region with the primer sets 515F(5'-GTGYCAGCMGCCGCGGTAA-3') / 806R(5'- GGACTACHVGGGTWTCTAAT-3') and ITS1FI2(5'-GTGARTCATCGAATCTTTG-3') / ITS2(5'-TCCTCCGCTTATTGATATGC-3'), respectively. PCR amplification was performed in a 50 μL mixture containing 12.5 μL of 2X Phanta Max master mix (Thermo Scientific), 2.5 μL of forward primer, 2.5 μL of reverse primer, 50 ng of DNA as a template, and 25 μL of sterile ddH2O. The PCR procedure for the bacterial 16S rRNA gene was as follows: initial denaturation at 98 °C for 30 s; 35 cycles of 98 °C for 10 s, 54 °C for 30 s, and 72 °C for 45 s; and a final extension at 72 °C for 10 min. For the fungal ITS2 region, the PCR procedure consisted of an initial denaturation at 98 °C for 30 s, 32 cycles of denaturation at 98 °C for 10 s, annealing at 54°C for 30 s, and extension at 72 °C for 45 s, and a final extension at 72 °C for 10 min. The PCR products were purified with AMPure XT beads (Beckman Coulter Genomics, Danvers, MA, USA) and quantified with a Qubit fluorometer (Invitrogen, USA). The amplicon pools were prepared for sequencing, and the size and quantity of the amplicon library were assessed on an Agilent 2100 Bioanalyzer (Agilent, USA) and with a Library Quantification Kit for Illumina (Kapa Biosciences, Woburn, MA, USA), respectively. The libraries were sequenced on a NovaSeq 6000 platform at LC-BIO Biotech Ltd. (Hangzhou, China). High-quality sequences were obtained after removal of low-quality sequences (quality score < 20 and sequence length < 100 bp). Chimeric sequences were filtered using Vsearch software (v2.3.4). After dereplication using DADA2, we obtained an amplicon sequence variant (ASV) feature table and feature sequence. The singleton ASVs were discarded. Taxonomic identification of bacteria and fungi was performed against the SILVA (v138) (Quast et al., 2013) and UNITE (v8.0) databases (Nilsson et al., 2018). Alpha diversity was calculated by QIIME2, where the same number of sequences was extracted randomly by reducing the number of sequences to the minimum of some samples. All the sequences obtained in this study were deposited in the National Center for Biotechnology Information (NCBI) GenBank under the SRA accession number PRJNA1034946, PRJNA1034808, PRJNA1182656 and PRJNA1185034.

Fungal mycelia DNA of 70 strains was also extracted using the cetyltrimethylammonium bromide (CTAB) method (Stewart and Via, 1993). We amplified the ITS region of the fungal DNA with the primers ITS4(5'-TCCTCCGCTTATTGATATGC-3') and ITS5(5'-GGAAGTAAAAGTCGTAACAAGG-3'). PCR was performed in a Veriti 96-well thermal cycler (Applied Biosystems, Inc., Foster City, CA, USA) in a 50 reactions volume composed of 25 μL of 2 × PCR Master Mix, 1 μL of each primer (10 μM), 22 μL of ddH2O and 1 μL of template DNA. The PCR procedure consisted of an initial denaturation at 94°C for 1 min; 35 cycles of denaturation at 94°C for 1 min, annealing at 54°C for 1 min, and extension at 72°C for 1 min; and a final extension at 72°C for 10 mins. PCR products were purified, and forward amplicons were sequenced by Sangon Biotech Co., Ltd. (Shanghai, China).

The obtained sequences were edited using EDITSEQ and SEQMAN software in the DNASTAR package (DnaStar, Inc., Madison, WI, USA). We aligned sequences in MEGA v. 6.0 using MUSCLE with default parameters (Edgar, 2004; Tamura et al., 2013), followed by manual checking of alignments. Taxonomic identification was performed via BLASTN analyses against the GenBank database. The ITS sequences reported in this study were deposited in the GenBank database (for accession numbers, see Table S2).

### References

**Edgar, R.C.** (2004). MUSCLE: multiple sequence alignment with high accuracy and high throughput. Nucleic Acids Res. **32:** 1792-1797.

**Nilsson, R.H., Larsson, K.-H., Taylor, A.F S., Bengtsson-Palme, J., Jeppesen, T.S., Schigel, D., Kennedy, P., Picard, K., Glöckner, F.O., Tedersoo, L., Saar, I., Kõljalg, U., and Abarenkov, K.** (2018). The UNITE database for molecular identification of fungi: handling dark taxa and parallel taxonomic classifications. Nucleic Acids Research **47:** D259-D264.

**Quast, C., Pruesse, E., Yilmaz, P., Gerken, J., Schweer, T., Yarza, P., Peplies, J., and Glockner, F.O.** (2013). The SILVA ribosomal RNA gene database project: improved data processing and web-based tools. Nucleic acids research **41:** D590-596.

**Stewart, C.N., and Via, L.E.** (1993). A rapid CTAB DNA isolation technique useful for rapid fingerprinting and other PCR applications. Biotechniques **14:** 748-750.

**Tamura, K., Stecher, G., Peterson, D., Filipski, A., and Kumar, S.** (2013). MEGA6: Molecular Evolutionary Genetics Analysis Version 6.0. Molecular Biology and Evolution **30:** 2725-2729.

### 2. Supplementary figures

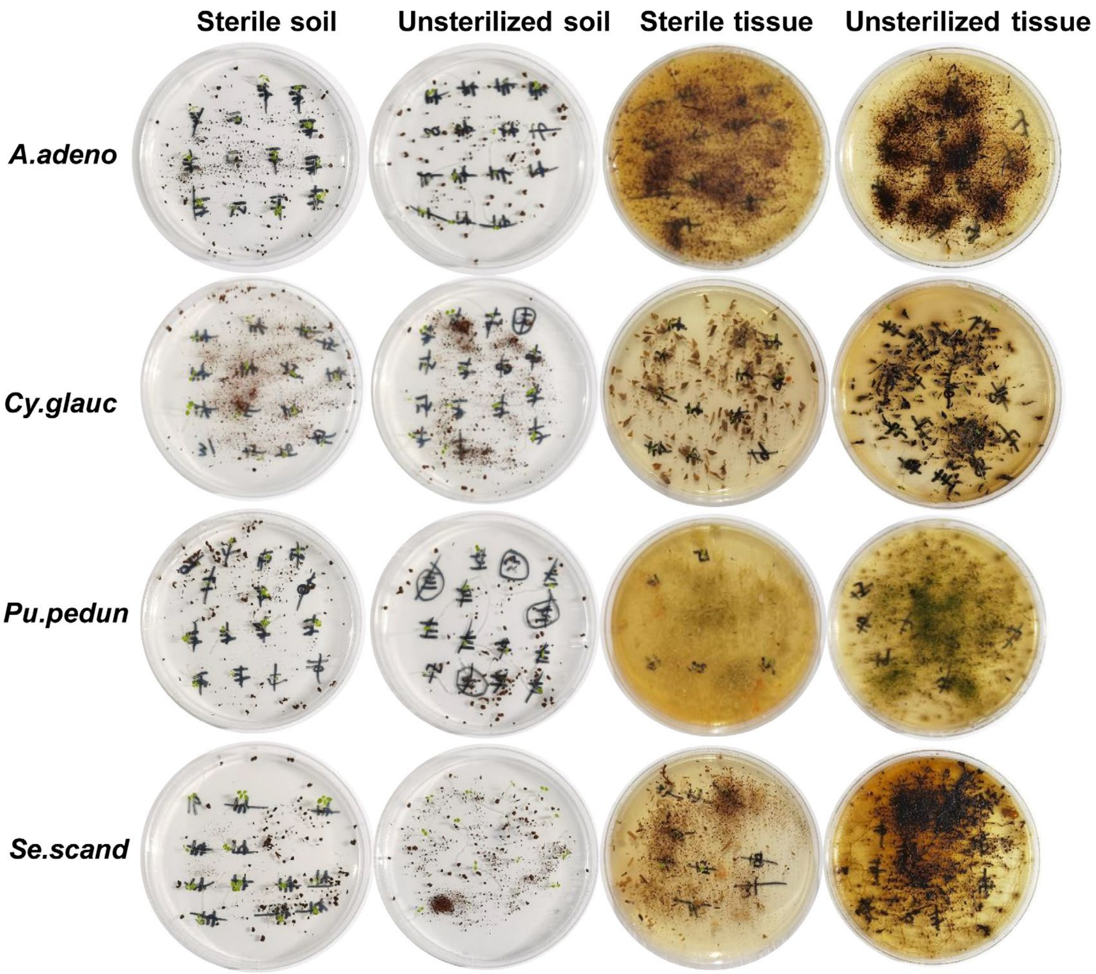


**Figure S1. Seedlings were inoculated with soils or plant tissues (from four randomly selected species) 21 days after sowing.** Only four randomly selected species are shown. *Ag. adeno*: *Ageratina adenophora*; *Cy.glauc*: *Cyclobalanopsis glaucoides; Pu.pedun*: *Pueraria peduncularis*; Se.scand: *Senecio scandens.*

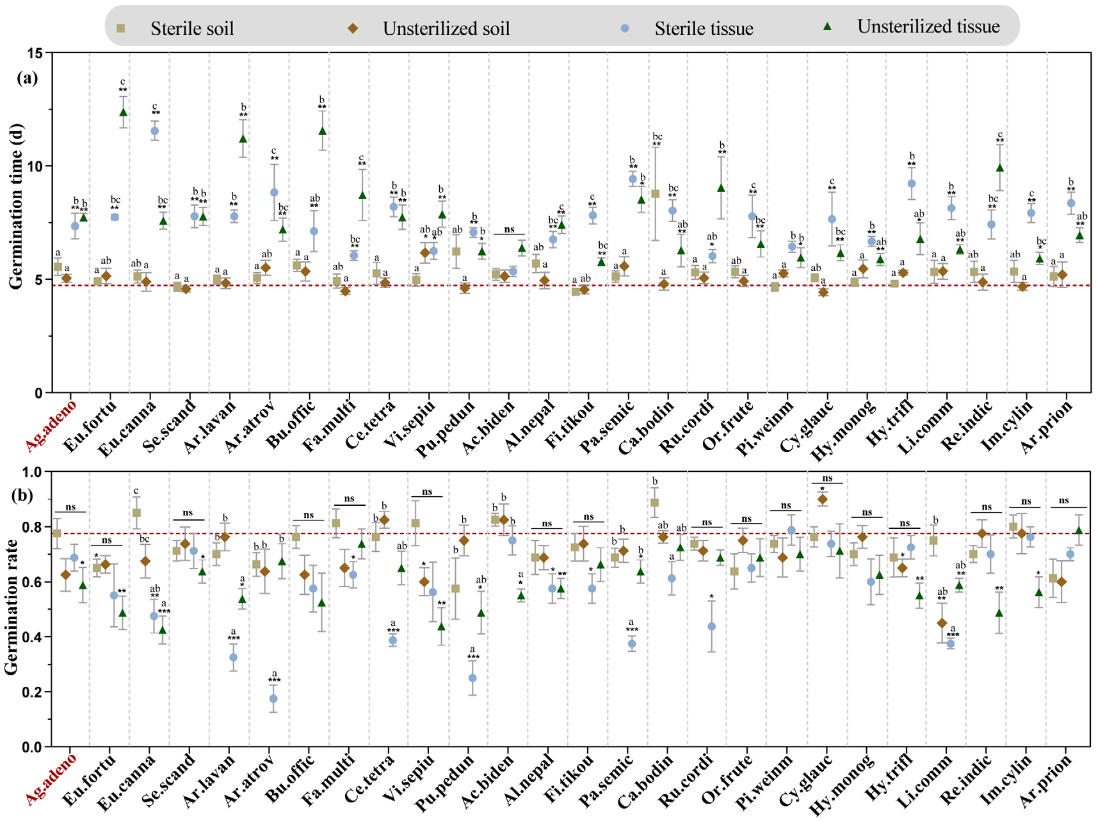


**Figure S2. Effects of sterile and unsterilized plant tissue or soil inoculation on *A. adenophora* germination time (a) and germination rate (b) among the 26 species.** Black * represents a significant difference compared with the control via Nonparametric Mann-Whitney U tests. Different lowercase means significant difference among four treatment groups of one species via Kruskal-Wallis test. The red dotted line represents the average value for the control treatment (no tissue or soil inoculated). Germination time of control: 4.73±0.211, germination rate of control: 0.775±0.042. The "ns" means nonsignificant. * $P < 0.05$, ** $P < 0.01$, *** $P < 0.001$. The abbreviations used for the abscissa are listed in Table S1, red text highlights exotic *A. adenophora* own tissue and soil inoculated. Means ± 1 SE, n=5.

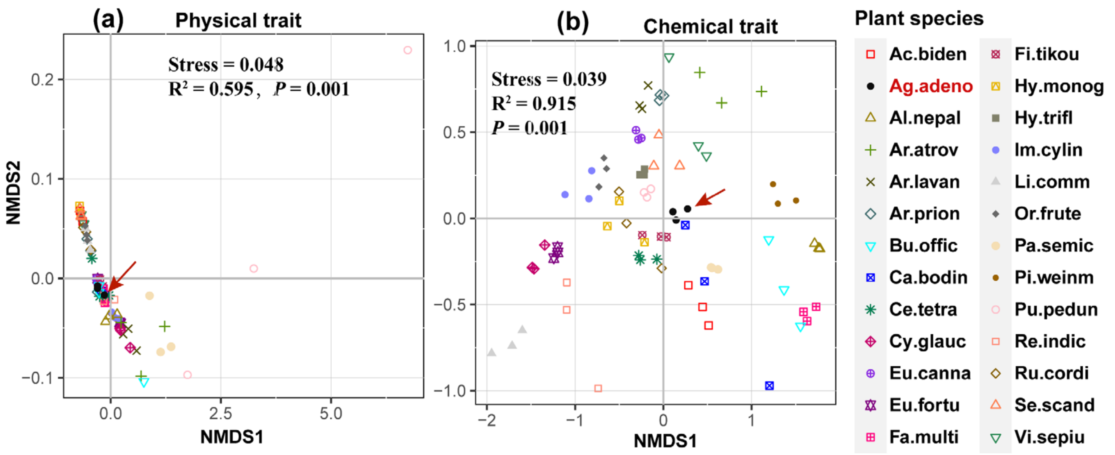


**Figure S3. Leaf physical (a) and chemical (b) trait dissimilarity for 26 plant species.** For plant species names, see Table S1, red arrows and text highlight the exotic species *A. adenophora*. $R^2$ means explained variation by plant species, $P$<0.05 represents significant dissimilarities among plant species.

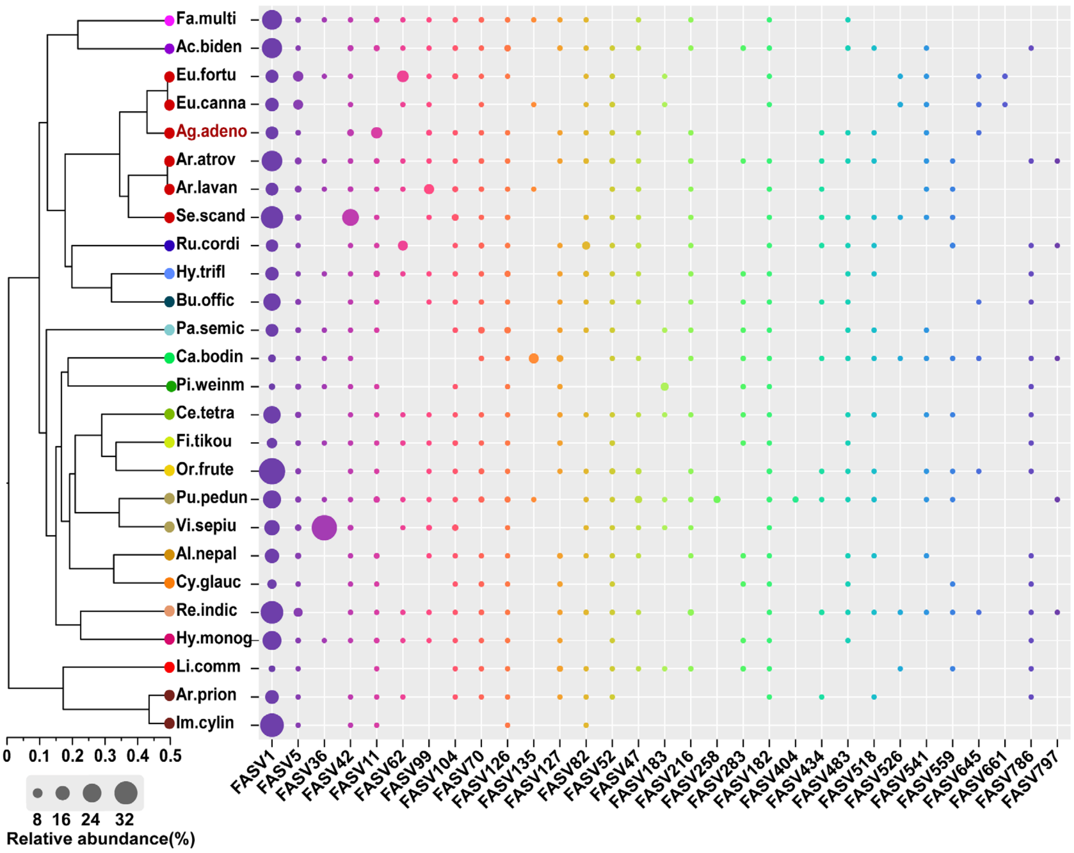


**Figure S4. The relative abundance of predictive fungal ASVs significantly correlated with seedling survival associated with aboveground tissues of 26 species.** The node colours in the phylogenetic tree represent the different plant families (Table S1). Red texts highlight the exotic species *A. adenophora*.
